# Fluency shaping increases integration of the command-to-execution and the auditory-to-motor pathways in persistent developmental stuttering

**DOI:** 10.1101/2020.07.27.219360

**Authors:** Alexandra Korzeczek, Annika Primaßin, Alexander Wolff von Gudenberg, Peter Dechent, Walter Paulus, Martin Sommer, Nicole E. Neef

## Abstract

Fluency-shaping enhances the speech fluency of persons who stutter, yet underlying conditions and neuroplasticity-related mechanisms are largely unknown. While speech production-related brain activity in stuttering is well studied, it is unclear whether therapy repairs networks of altered sensorimotor integration, imprecise neural timing and sequencing, faulty error monitoring, or insufficient speech planning. Here, we tested the impact of one-year fluency-shaping therapy on resting-state fMRI connectivity within sets of brain regions subserving these speech functions. We analyzed resting-state data of 22 patients who participated in a fluency-shaping program, 18 patients not participating in therapy, and 28 fluent control participants, measured one year apart. Improved fluency was accompanied by an increased synchronization within the sensorimotor integration network. Specifically, two connections were strengthened; the left inferior frontal gyrus showed increased connectivity with the precentral gyrus at the representation of the left laryngeal motor cortex, and the left inferior frontal gyrus showed increased connectivity with the right superior temporal gyrus. Thus, therapy-associated neural remediation was based on a strengthened integration of the command-to-execution pathway together with an increased auditory-to-motor coupling. Since we investigated task-free brain activity, we assume that our findings are not biased to network activity involved in compensation but represent long-term focal neuroplasticity effects.

## Introduction

Most people speak fluently with ease, but fluent speech requires a complex interplay across multiple functional domains. In stuttering, aberrant brain activity and connectivity is evident in networks that convey fluent speech production (Ingham et al., 2018). However, especially in adults who experienced lifelong stuttering, it is challenging to differentiate core neural deficits from stuttering-induced neural signatures or intervention-induced neuroplasticity from compensatory network activity (Kell, 2012). Thus, our understanding of the neurophysiological mechanistic principles of stuttering and its neural remediation remains limited.

The neural networks that may become involved during stuttering intervention have hardly been studied. Previous neuroimaging studies reported local and distributed intervention-induced activity changes in the left inferior frontal gyrus (Kell et al., 2009; Lu et al., 2017; Neumann et al., 2018), cerebellum (De Nil et al., 2001; Lu et al., 2012; Toyomura et al., 2015), and basal ganglia (Toyomura et al., 2015), and a change in lateralization of speech-related frontal brain activity towards the more typical leftwards pattern (De Nil et al., 2003; Kell et al., 2009; Neumann et al., 2018, 2003). All except one earlier study (Toyomura et al., 2015) involved the training of voicing patterns to shape speech fluency, a common approach to overcome stuttering.

Fluency shaping is a speech restructuring method that requires individuals to change their speech patterns. Specifically, during fluency shaping, patients learn to speak slowly with gentle onsets of phonation, light articulatory contacts, and soft voicing of plosives (Euler et al., 2009; Webster, 1974). After three weeks of fluency shaping training, connectivity was increased between the left anterior superior temporal gyrus and the left articulatory motor cortex, and hyperconnectivity was reduced between the left IFG pars opercularis and the sensory feedback processing left supramarginal gyrus (Kell et al., 2018). This observation suggests a treatment-induced boost of auditory-to-motor coupling and likely indicates neuroplasticity induced by sensorimotor learning (Calmels, 2020). However, a stuttering intervention that leads to a significant reduction of speech dysfluencies in adults (Euler et al., 2009) and children (Euler et al., 2021) presumably addresses large-scale functional networks that exceed auditory-to-motor mapping. Further supporting domains that are related to speech processing could be speech planning (Andreatta et al., 2010; Price, 2012), sensorimotor integration (Behroozmand et al., 2015; Darainy et al., 2019; Hickok et al., 2011; Tourville et al., 2008), articulatory convergence (Brown et al., 2005; Guenther, 2016; Turkeltaub et al., 2002), and the inhibition of competitive processes (Ghahremani et al., 2018; Xue et al., 2008). Previous studies might have failed to detect changes in these crucial functional domains because measured brain activity was biased by the tasks employed.

One suitable approach to scrutinizing learning-induced neuroplasticity is resting-state functional magnetic resonance imaging (rs-MRI). On the one hand, rs-fMRI is free from confounds of task performance, particularly in participants who may present symptoms such as physical concomitants during speaking. Thus, task-free brain activity assesses changes in brain dynamics that are not biased by differences in how a task is performed in pre-learning versus post-learning condition (Vahdat et al., 2011). On the other hand, it is widely assumed that ongoing spontaneous global activity of the brain at rest is (1) highly-structured, (2) closely relates to underlying anatomical connectivity, and (3) reflects local neuronal dynamics, signal transmission delay, and genuine noise, i.e., unstructured input (Deco et al., 2011). It has been shown that even under the resting-state condition, brain areas show activity changes with learning, and correlated activity increases between learning-related areas (Albert et al., 2009; Darainy et al., 2019; Vahdat et al., 2011).

To date, task-free brain activity has been studied twice to test stuttering intervention-induced neuroplasticity (Lu et al., 2017, 2012). Both studies investigated the same speech therapy. During the 7-day intervention with three daily sessions, participants trained a new voicing pattern with word listen-and-repeat tasks followed by overt-Pinyin-reading tasks. Later, participants listened to their audio-recordings and received the therapist’s feedback. In addition, participants applied the newly learned speaking pattern to utterances produced throughout their daily lives (Lu et al., 2017). The first study showed an intervention-related decrease of rs-fMRI connectivity in the left declive and vermis area of the cerebellum. In addition, connectivity changes in the cerebellum correlated positively with the change of stuttering severity after the change of duration of the stuttering events and physical concomitants was regressed out (Lu et al., 2012). Because stuttering-related cerebellar overactivity was often considered to reflect compensatory activity (Brown et al., 2005; De Nil et al., 2008; Lu et al., 2010; Watkins et al., 2008), Lu and colleagues suggested the reduced cerebellar rs-fMRI connectivity to display reduced compensatory activity and, thus, to indicate a neural reorganization of the intrinsic functional architecture of speech processing. The second study, which included a reduced number of the same participants, showed that rest-related connectivity changes in the cerebellum and task-related activity increases in the left ventral inferior frontal gyrus and insula, were not correlated. The task-related activity was measured during the overt reading of monosyllabic Chinese characters (Lu et al., 2017). This observation was discussed to support the idea that resting-state functional connectivity and task-related brain activity provide different insights into mechanisms behind brain plasticity.

Here, we use a longitudinal approach to examine stuttering intervention-induced improvement in speech fluency and neurofunctional reorganization. To this end, we acquired rs-fMRI data before and 11 months after a computer-assisted fluency shaping training (Euler et al., 2009) in persons with developmental stuttering (PDS+) and tested time-dependent connectivity changes. We controlled for the specificity of intervention-induced changes by studying two control groups, i.e., patients with developmental stuttering not taking part in any stuttering intervention (PDS−) and fluent controls (FC). We quantified the synchronicity of spontaneous low-frequency fluctuations to characterize the connectivity between functionally related brain hubs. We determined sets of ROIs to assess connectivity of a priori determined semi-discrete networks associated with speech planning (Bohland and Guenther, 2006; Rottschy et al., 2012), articulatory convergence (Guenther, 2016), speech-related sensorimotor integration (Darainy et al., 2019), and speech motor inhibitory control (Ghahremani et al., 2018; Neef et al., 2016). All ROIs were chosen from the literature. However, we made no directional hypothesis before the data collection. Chosen ROIs capture spontaneous BOLD fMRI fluctuations of neuronal populations that are integral parts of brain networks. Network dynamics are without much doubt nonlinear and difficult to predict. In this vein, the outcomes were not predicted and are therefore exploratory. To capture brain-behavior relationships, we additionally explored correlations between changes in functional connectivity and speech fluency.

## Methods

### Participants

The current data were collected during a dissertation project (Primaßin, 2019) that evaluated the long-term effects of an intensive stuttering intervention on white matter integrity and task-related brain activity. Persons who stutter who were about to begin with intervention at the Kassel Stuttering Therapy (KST) were invited to participate in the MRI study. Volunteers from the KST were assigned to PDS+. Fluent control participants were recruited via advertisements at the homepage of the Department of Clinical Neurophysiology and at notice boards of the university campus and clinic and were assigned to FC. Stuttering controls were recruited via announcements at stuttering self-help groups and at the 2016 annual congress of the German stuttering self-help group association (BVSS). Stuttering controls did not participate in any stuttering intervention during the entire study period and were assigned to PDS−.

Seventy-six right-handed, monolingual speakers of German participated voluntarily in the current study. Exclusion criteria were speech or language disorders other than developmental stuttering, neurological impairment, drug abuse, or medications that act on the central nervous system. None of the PDS− took part in any stuttering intervention during the entire study period. For analysis, we excluded the data of three PDS+ because they participated in addition in a different stuttering intervention. Data of further four participants (1 PDS+, 2 PDS−, 1 FC) were excluded because of missing behavioral or rs-fMRI data, and data of one PDS+ were excluded because of extensive rs-fMRI motion artifacts. Thus, the rs-fMRI data analysis comprised 22 PDS+ (2 females, mean age 25.6 ± 11.7 years with 7 participants younger than 18 years), 18 PDS− (2 females, mean age 34.8 ± 7.0 years, with no participant younger than 18 years), and 28 FC (4 females, mean age 25.1 ± 7.4 years with 5 participants younger than 18 years). While age was comparable between PDS+ and FC with *T* = −0.16 and *p* = 0.87, PDS+ were younger than PDS− with *T* = −3.1 and *p* = 0.004, and PDS+ were younger than PDS− with *T* = −4.49 and p < 0.001. PDS+ and FC were matched with regard to sex and handedness (Oldfield, 1971). PDS− had a higher education score than participants in the two other groups (see Table 1). Speech fluency (Stuttering severity index, SSI-4) (Riley et al., 2004) of all participants was assessed prior to each MRI session. Stuttering severity was lower in PDS− than in PDS+ (Table 1). In addition, a self-assessment of the psychosocial impact of stuttering (Overall Assessment of the Speaker’s Experience of Stuttering, OASES) (Yaruss and Quesal, 2014) indicated that PDS+ were more affected by stuttering than PDS−. Finally, both PDS− and PDS+ were comparable regarding the time span in years that had passed since the last stuttering intervention before participating in the current study (Table 1 and Supplementary Table 1). There were three participants per group not providing information on their stuttering intervention history. Age of intervention and nature of intervention varied in both groups. However, the groups were too small to compare them with respect to these two variables.

**Table 1.** Demographic data of participants.

|  | PDS+ | PDS- | FC | Test-statistics<br>(*df*) | two-sided<br>*p*-value |
| --- | --- | --- | --- | --- | --- |
| *n* | 22 | 18 | 28 |  |  |
| Age, years | $25.6 \pm 11.7$ | $34.8 \pm 7.0^*$ | $25.1 \pm 7.4$ | $7.58 (2, 65)^i$ | 0.001 |
| Sex ratio | 20:2 | 16:2 | 24:4 | —<sup>ii</sup> | 0.89 |
| Education<sup>a</sup> | 2 (1.0)<sup>-</sup> | 6 (3.0)<sup>*</sup> | 3 (2.8) | $16.68 (2, 68)^{iii}$ | < 0.001 |
| Handedness | 91 (12) | 91 (33) | 100 (33) | 0.04 (2,68)<sup>iii</sup> | 0.98 |
| SSI-4 at T1 | 25 (14.3)<sup>#</sup> | 14 (11.3) | — | 2.56<sup>iv</sup> | 0.010 |
| SSI-4 at T2 | 9 (10.5) | 12.5 (11.0) | — | -1.31<sup>iv</sup> | 0.194 |
| OASES at T1 | 3.0 (0.6)<sup>#</sup> | 2.0 (0.4) | — | 4.70<sup>iv</sup> | < 0.001 |
| OASES at T2 | 1.9 (0.5) | 2.0 (0.5) | — | -0.65<sup>iv</sup> | 0.516 |
| Last therapy, years ago | 8.5 ± 8.7<sup>~</sup> | 12.7 ± 8.8<sup>~</sup> |  | -1.18(32)<sup>v</sup> | 0.249 |
| Onset, years | 4.8 ± 3.0 | 5.0 ± 3.6 | — | 0.22<sup>iv</sup> | 0.839 |
| MRI interval, months | 11.6 ± 1.0 | 11.6 ± 1.4 | 11.4 ± 0.8 | 0.95 (2)<sup>iii</sup> | 0.623 |

Interval/ratio -scaled variables are presented as mean ± standard deviation. Ordinal-scaled variables are presented as median (interquartile range). \*significantly different from both other groups in post hoc comparisons ( $p < 0.001$ ), <sup>#</sup>significantly different from stuttering controls ( $p < 0.001$ ), <sup>~</sup> three missing values, <sup>i</sup>one-way independent ANOVA, <sup>ii</sup>Fisher's exact test, <sup>iii</sup>Kruskal-Wallis test, <sup>iv</sup>Mann-Whitney test, <sup>v</sup>unpaired t-test, <sup>a</sup>achieved education levels were 1 = still attending school, 2 = school, 3 = high school, 4 = <2years college, 5 = 2 years of college, 6 = 4 years of college, 7 = postgraduate

The study was registered on January 14, 2016, at the study center of the University Medical Center of Göttingen and was given the registration number 01703. This registration is not publicly accessible, but access could be requested under the following email address. The ethical review board of the University Medical Center Göttingen, Georg August University Göttingen, Germany, approved the study, and all participants gave their written informed consent, according to the Declaration of Helsinki, before participation. In addition, informed consent was obtained from parents or legal guardians of participants under the age of 18.

All participants took part in two MRI sessions (T1 and T2) separated by 10 to 15 months. The scanning interval was similar between groups (Table 1). PDS+ were scanned pre- T1) and post-intervention (T2).

### Intensive stuttering intervention and follow-up care

PDS+ took part in the Kasseler Stottertherapie (Euler et al., 2009), an intensive program that incorporates fluency shaping with computer-assisted biofeedback during a two-week on-site and one-year follow-up treatment. Fluency shaping reconstructs patterns of vocalization, articulation, and respiration, resulting in prolonged speech, soft voice onsets of initial phonemes, and a smooth transition between sounds. It was first introduced with the precision fluency shaping program by Webster (Max Ludo and Caruso Anthony J., 1997; Webster, 1980, 1974). The overarching aim of this approach is to train to speak slowly with gentle onsets of phonation, light articulatory contacts, and soft voicing of plosives. In the current study, the on-site intervention encompassed two weeks of intensive therapy and training, i.e., at least eight hours per day and seven days per week. The intervention was structured into alternating sessions. Sessions included group therapy, individual computer-assisted speech training, one-to-one speech therapy, and in-vivo training. In-vivo training stands for applying the speech technique in real-life situations that require patients to talk to persons outside the therapy setting while still receiving support from a therapist. Next to applying the speech technique during everyday communication, participants were encouraged to practice daily with the computer. Computer-assisted training at home was mainly based on the biofeedback-assisted practice of the new speech patterns. Biofeedback consisted of a visualization of the speech sound wave. As an incentive, participants could get the costs of the software reimbursed by their health insurance if they practiced at least 1980 minutes within the first half of the year and 990 minutes within the second half (Euler et al., 2009). Thus, the intervention under study was the same as in Kell et al. (2018, 2009) and Neumann et al. (2018, 2005, 2003), differed in the way of providing feedback from the one in De Nil et al. (2003) and differed in intervention duration, therapy content, and provision of feedback from the stuttering intervention studied in the only other resting-state study (Lu et al., 2017, 2012). During the follow-up period, there were two refresher courses at the therapy center at one month and ten months, respectively, after the initial intensive training. If participants were not able to attend the 10-month refresher, they could also attend subsequently offered refresher courses. In this study, participants scheduled the second refresher at the latest 14 months after the intervention. On rare occasions, due to organizational issues, MRI measurements at T2 took place one day before the second refresher.

### Assessment and statistical analysis of behavioral data

Changes in speech fluency were assessed by two experienced speech-language pathologists (one of whom was A.P.) using the Stuttering Severity Index (SSI-4). The SSI-4 assessment included a spontaneous speech sample and a reading sample. Each sample comprised 488 to 500 syllables. For interrater reliability estimation, the two raters analyzed nine randomly chosen participants, three from each group. Reliability estimates were statistically assessed with SPSS software with Krippendorf’s Alpha Reliability Estimate (KALPHA) using 10,000 bootstrapping samples at an ordinal level (Hayes and Krippendorff, 2017). KALPHA ranged between 0.84 and 0.98 for the SSI-4 sub-scores reading, spontaneous speech, duration, and concomitants. KALPHA was 0.96 for the SSI-4 total score, indicating a good to excellent consensus between raters. The participants’ experience with stuttering was assessed with the German version of the OASES (Yaruss and Quesal, 2014). We assessed behavioral changes as a change in the SSI-4 total scores and change in the OASES total scores between T1 and T2 using R (version 3.5.3). We ran robust mixed ANOVAs on trimmed means with Group as between-factor and Time as within-factor using the function tsplit with the default trimming level of 0.2 of the package WRS (R.R. Wilcox’ robust statistics functions -version 0.37). Time was implemented as the second factor. Post hoc we applied Wilcoxon signed-rank tests.

### Definition of four speech-related semi-discrete brain networks

Resting-state fMRI captures brain activity in the absence of a task. Spontaneous fluctuations of the blood-oxygen-level-dependent (BOLD) signal represent specific patterns of synchronous activity and reflect the functional organization of the brain (Biswal et al., 1995). Compared to data-driven approaches, which are also common to study rs-fMRI activity, ROI-to-ROI analyses provide detailed information on the specific connectivity of brain areas of interest as demonstrated for the dorsal and ventral attentional systems (Fox et al., 2006) or the functional connectivity of the anterior cingulate cortex (Margulies et al., 2007). Fluent speech production engages large-scale brain networks conveying emotional, linguistic, cognitive, sensory, and motor functions. Among these processes, dysfunctional speech planning, articulatory convergence, sensorimotor integration or motor inhibition most likely cause the primary motor signs of stuttering, which are sound and syllable repetitions, sound prolongations, and speech blocks. Here, we distinguished four semi-discrete brain networks consisting of brain regions that are recruited for any of these functions (Fig. 1).

**Figure 1.**
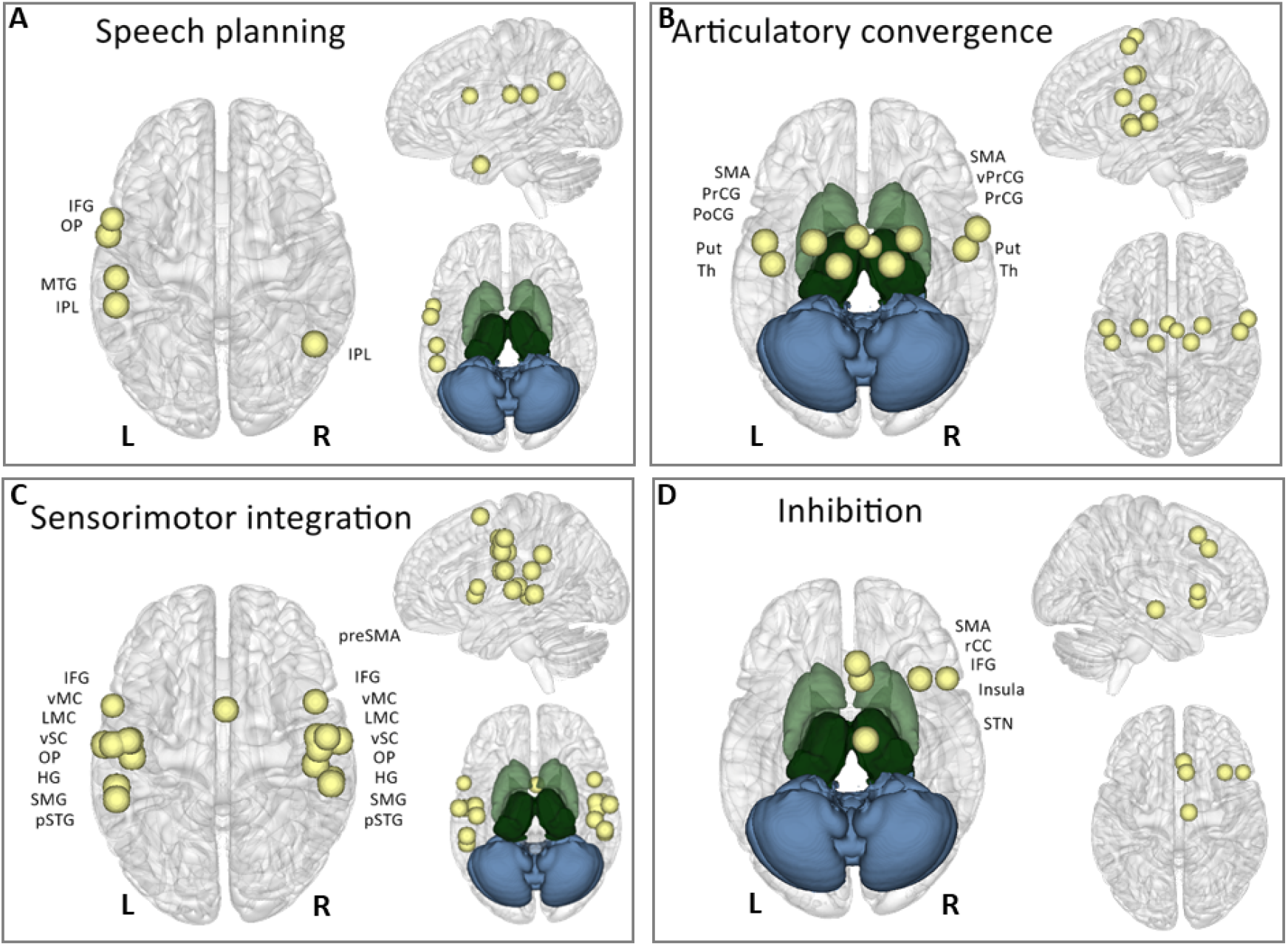
ROI-to-ROI resting state fMRI analyses were conducted in four semi-discrete functional networks. Spheres with a diameter of 6 mm served as seed regions, overlaid here on rendered surfaces of the MNI standard brain. Intervention effects were tested for speech planning (A), articulatory convergence (B), sensorimotor integration (C), and motor inhibition (D). HG = Heschl’ s gyrus, primary auditory cortex; IPL = inferior parietal lobe; IFG = inferior frontal gyrus, pars opercularis; LMC = laryngeal motor cortex; MTG = middle temporal gyrus; O*p* = parietal operculum; PoCG = postcentral gyrus; PrCG = precentral gyrus; preSMA = pre-supplementary motor area; pSTG = posterior superior temporal gyrus; Put = putamen; rCC = rostral cingulate zone; SMA = supplementary motor area; SMG = supramarginal gyrus; STN = subthalamic nucleus; Th = thalamus; vMC = ventral primary motor cortex; vPrCG = ventral precentral gyrus; vSC = ventral primary somatosensory cortex.

Several left-hemispheric regions contribute to speech planning: posterior inferior gyrus, insula, temporoparietal regions, and the proper and pre-supplementary motor area (Andreatta et al., 2010; Price, 2012). A selected brain region of dysfunctional speech planning was derived from a fixed-effects analysis of an earlier fMRI study of our lab that investigated imagined speaking compared with humming in 15 PDS and 15 FC (Neef et al., 2016) (Fig. 1A). Between-group contrasts of an ROI analysis within the area BA 44 and of a functional connectivity analysis with the left posterior area 44 as seed revealed dysfunctional brain regions of speech planning in PDS (Neef et al., 2016). Articulatory convergence seeds originated from combined ALE meta-analyses (Guenther, 2016) on brain imaging studies of simple articulatory movements of the jaw, larynx, lips, tongue, and respiratory system (Fig. 1B). The rationale behind this concept was that speaking requires the joint coordination of multiple articulatory subsystems, i.e., jaw movements, lip movements, larynx movements, respiratory movements, and tongue movements. Brain regions that are involved in the control of multiple articulatory subsystems were defined as regions of high articulatory convergence. To determine such brain regions, Guenther (2016) performed five activation likelihood estimate (ALE) meta-analyses, one for each subsystem, thereby including only functional imaging contrasts of non-speech movement tasks. Afterward, he identified brain coordinates where foci of three or more articulatory systems showed very close proximity by visual inspection(Guenther, 2016). Speech-related sensorimotor integration seeds were derived from a ‘listen-and-repeat’ localizer task in a brain imaging study of sensorimotor plasticity in speech motor adaptation (Darainy et al., 2019) (Fig. 1C). The study investigated in 19 neurotypical participants on two consecutive days whether behavioral learning-related changes in perception and speech movements influenced brain motor areas directly or indirectly via sensory areas. ROI-to-ROI analyses of rs-functional connectivity were used to identify sensorimotor plasticity between the first and the second day (Darainy et al., 2019). Of note, the ROI coordinates of the primary motor cortex (mid precentral gyrus) reported by (Darainy et al., 2019) were exchanged because of the differences in speech tasks. Whereas participants in Darainy et al. (2019) adjusted articulatory movements to the perception of vowels, the stuttering intervention required the modification of voicing and thus a skillful control of the larynx. (Neumann et al., 2018) for example reported that the very same stuttering intervention normalized the mean fundamental frequency (meanF0) for PWS+. Thus, coordinates of the laryngeal motor cortex (LMC) that were chosen for replacement, were derived from a fMRI meta-analysis encompassing 19 overt speech production studies with 283 neurotypical participants (Kumar et al., 2016; Simonyan, 2014). For further rs-fMRI analyses in CONN (SPM toolbox), the coordinates in Talairach space, left LMC at [− 45, −14, 33] and right LMC at [44, −12, 35], reported by Kumar et al. (2016), were converted with GingerALE (http://www.brainmap.org/ale/) using the transform “Talairach to MNI (SPM)” to MNI space, left LMC [−47, −10, 34] and right LMC [49, −8, 35]. Motor inhibition seeds that involved common areas of inference resolution, action withholding, and action cancellation were derived from a meta-analysis of 225 studies (Zhang et al., 2017). We added the subthalamic nucleus seed to the inhibition network to account for the dedicated involvement of this structure in response inhibition (Aron and Poldrack, 2006) (Fig. 1D). In two experiments, this study showed the inhibitory role of the subthalamic nucleus using action cancellation tasks (stop-signal task)(Aron and Poldrack, 2006). We created spherical seeds with a radius of 6 mm for all ROIs. Coordinates for brain hubs involved in speech-related sensorimotor integration can be found in Table 2. Seeds for the remaining three networks are listed in Supplementary Tables 3-5. Seed ROIs did not overlap.

**Table 2.**
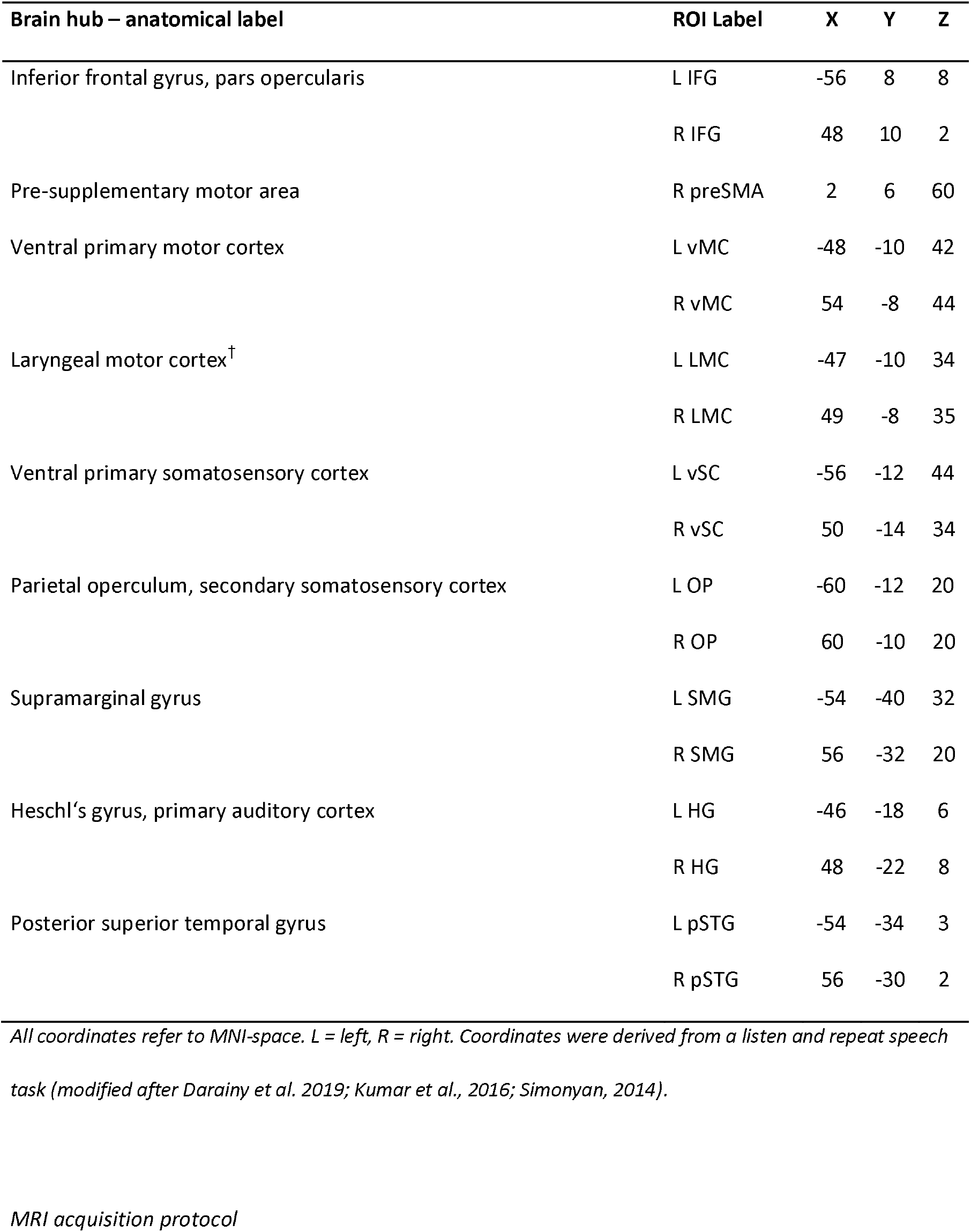
Brain hubs of speech-related sensorimotor integration.

### MRI acquisition protocol

MRI data were acquired in a 3 Tesla Siemens Magnetom Tim Trio scanner (Erlangen, Germany) using an eight-channel, phased-array head coil at the University Medical Center Göttingen, Germany. Sagittal T1-weighted structural data were acquired with a 3D turbo fast low angle shot (FLASH) sequence (TR = 2250ms, TE = 3.26ms, TI = 900ms, flip angle = 9°, 256mm FoV, 7/8 Fourier phase encoding) as whole-brain anatomical reference data at a spatial resolution of 1 × 1 × 1 mm³ voxel size (256 × 256 matrix). For resting-state fMRI a gradient-echo echo-planar imaging (EPI) sequence (TR = 1800ms, TE = 30ms, flip angle = 70°, parallel acquisition factor 2, 192 mm FoV, 33 slices, 194 volumes) was used with isotropic voxels at 3 (mm)³ and a 64 x 64 acquisition matrix. We acquired two six-minute rs-fMRI time series at T1 and at T2, respectively, while participants fixated on a cross in an open eyes condition. Due to different head sizes, the rs-fMRI data did not fully cover the cerebellum in some participants. Therefore, the cerebellum was excluded in further rs-fMRI analyses. Participants lay in a supine position in the scanner and wore headphones for noise protection and MR-compatible LCD goggles (VisuaStim XGA, Resonance Technology Inc., Northridge, CA, USA).

### Rs-fMRI data preprocessing

Structural and functional MRI data were preprocessed and analyzed with CONN functional connectivity toolbox version 18b (Whitfield-Gabrieli and Nieto-Castanon, 2012). The toolbox is based on Matlab and Statistical Parametric Mapping (SPM). The standard preprocessing pipeline of CONN was used with functional realignment, functional centering of the image to (0, 0, 0) coordinates, slice-timing correction, structural centering to (0, 0, 0) coordinates, structural segmentation and normalization to MNI space, and spatial smoothing with a smoothing kernel of 8 mm full-width at half-maximum. Motion parameters and signal outliers were detected via the Artifact Rejection Toolbox set to 95^th^ percentile, which allowed for quantifying participant motion in the scanner and identifying outliers based on the mean signal (Goto et al., 2016; Power et al., 2012). Motion parameters, and white matter and cerebral spinal fluid signals were included as confounds and regressed out during denoising before first-level analysis. Data were denoised using a bandpass filter of 0.009 - 0.08 Hz.

### ROI-to-ROI functional connectivity analyses

We used the CONN toolbox to analyze rs-fMRI data to determine Fisher-transformed correlation coefficients of bivariate ROI-to-ROI correlations with hemodynamic response function weighting for each of the four sets of ROIs (Fig. 1). We then run four global mixed models ANOVAs with Group (PDS+, FC) as the between-subjects factor and Time and ROIs as within-subjects factors to test intervention-induced neuroplasticity. For multiple comparison correction, we set the connection threshold at seed level p-FDR < 0.05 (two-sided) and the seed-level permutation analysis threshold (based on a Network-Based Statistics by intensity approach^12^) at p-FDR < 0.05. Beta values represent average functional connectivity (Fisher-transformed correlation coefficients) and indicate effect sizes. If Group x Time interactions were significant, we extracted beta values and calculated two-sided paired t-tests in CONN to compare connectivity between T2 and T1 separately for each group. Furthermore, we calculated two-sided unpaired t-tests to analyze group differences separately at T1 and T2 (Table 3).

**Table 3.** Disentangling the Group × Time interaction with PDS+ and FC.

| Seed - target region | Time | Beta | T (DF) | p | Group | T (DF) | p |
| --- | --- | --- | --- | --- | --- | --- | --- |
|  | T2 > T1 |  |  |  | FC vs PDS+ |  |  |
| left IFG – left LMC | PDS+ | 0.1 | 4.91<sup>#</sup> (21) | < 0.001 | T1 | 1.84 (47.7) | 0.072 |
|  | FC | -0.051 | -1.81<sup>#</sup> (27) | 0.082 | T2 | -2.8 (35.9) | 0.007 |
| left IFG – right pSTG | PDS+ | 0.094 | 3.32<sup>#</sup> (21) | 0.003 | T1 | 1.38 (46.7) | 0.174 |
|  | FC | -0.051 | -1.46<sup>#</sup> (27) | 0.155 | T2 | -3.09 (48.0) | 0.003 |

<sup>#</sup>Positive values indicate increased connectivity, and negative values indicate decreased connectivity at T2.

PDS− were not included in the global ANOVAs because of the age differences. Still, to test whether the condition to continue to stutter for the period of the treatment influenced brain connectivity, we calculated in CONN additional two-sided independent t-tests, comparing the adjusted connectivity change from T1 to T2 (left IFG-to-LMC, left IFG-to-right pSTG) between PDS+ and PDS. Connectivity changes were adjusted for age and SSI total scores at T1. Because education correlated with age, r = 0.483, p < 0.001, education was not regressed out. Post-hoc we calculated two-sided paired t-tests to compare adjusted connectivity between T2 and T1 separately for each group. Furthermore, we calculated two-sided unpaired t-tests to analyze group differences separately at T1 and T2 (Table 4). Finally, Pearson correlations were calculated in MATLAB (R2018b) to test brain-behavior relationships.

**Table 4.** Disentangling the Group × Time interaction with PDS+ and PDS−.

| Seed - target region | Time | Beta | T (DF) | p | Group | T (DF) | p |
| --- | --- | --- | --- | --- | --- | --- | --- |
|  | T2 > T1 |  |  |  | PDS+ vs. PDS- |  |  |
| <sup>1</sup>left IFG – left LMC | PDS+ | 0.09 | 3.64<sup>#</sup> (19) | 0.002 | T1 | -1.34 (37.9) | 0.188 |
|  | PDS- | -0.08 | -1.73<sup>#</sup> (15) | 0.105 | T2 | 1.45 (36.9) | 0.155 |
| <sup>1</sup>left IFG – right pSTG | PDS+ | 0.11 | 3.26<sup>#</sup> (19) | 0.004 | T1 | -0.78 (34.2) | 0.442 |
|  | PDS- | 0.008 | 0.14<sup>#</sup> (15) | 0.891 | T2 | 1.69 (35.0) | 0.100 |

<sup>#</sup>Positive values indicate increased connectivity, and negative values indicate decreased connectivity at T2.

### Data availability statement

The data that support the findings of this study are available from the corresponding author upon reasonable request.

## Results

### Computer-assisted intensive intervention improved speech fluency and well-being

The observed reduction in the total scores of the Stuttering Severity Index (SSI-4) and the Overall Assessment of the Speaker’s Experience of Stuttering (OASES) indicates a positive effect of stuttering intervention. The robust ANOVA for stuttering severity revealed a significant interaction of Group by Time, Q = 24.44, p < 0.001, an effect of Group Q = 48.38, p < 0.001, and an effect of Time Q = 50.99, p < 0.001. Post hoc tests showed that stuttering severity decreased from T1 to T2, V = 253, p < 0.001, r = −0.87 (Fig. 2C) in the intervention group. In the non-intervention groups, on the other hand, the SSI-4 scores remained unchanged with V = 63, *p* = 0.81, r = −0.06 for PDS− and V = 58.5, *p* = 0.38, r = −0.17 for FC. Similarly, the robust ANOVA for the speaker’s experience of stuttering revealed a significant interaction of Group by Time, Q = 66.73, p < 0.001, an effect of Group Q = 20.39, p < 0.001 and an effect of time Q = 74.17, p < 0.001. Post hoc tests revealed a decrease in the OASES-scores between T1 and T2 only in PDS+, V = 253, p < 0.001, r = −0.87 (Fig. 2B). PDS− experienced no changes in their experience with stuttering, V = 110.5, *p* = 0.29, r = −0.25. Behavioral outcome measures are summarized in Supplementary Table 6.

**Figure 2.**
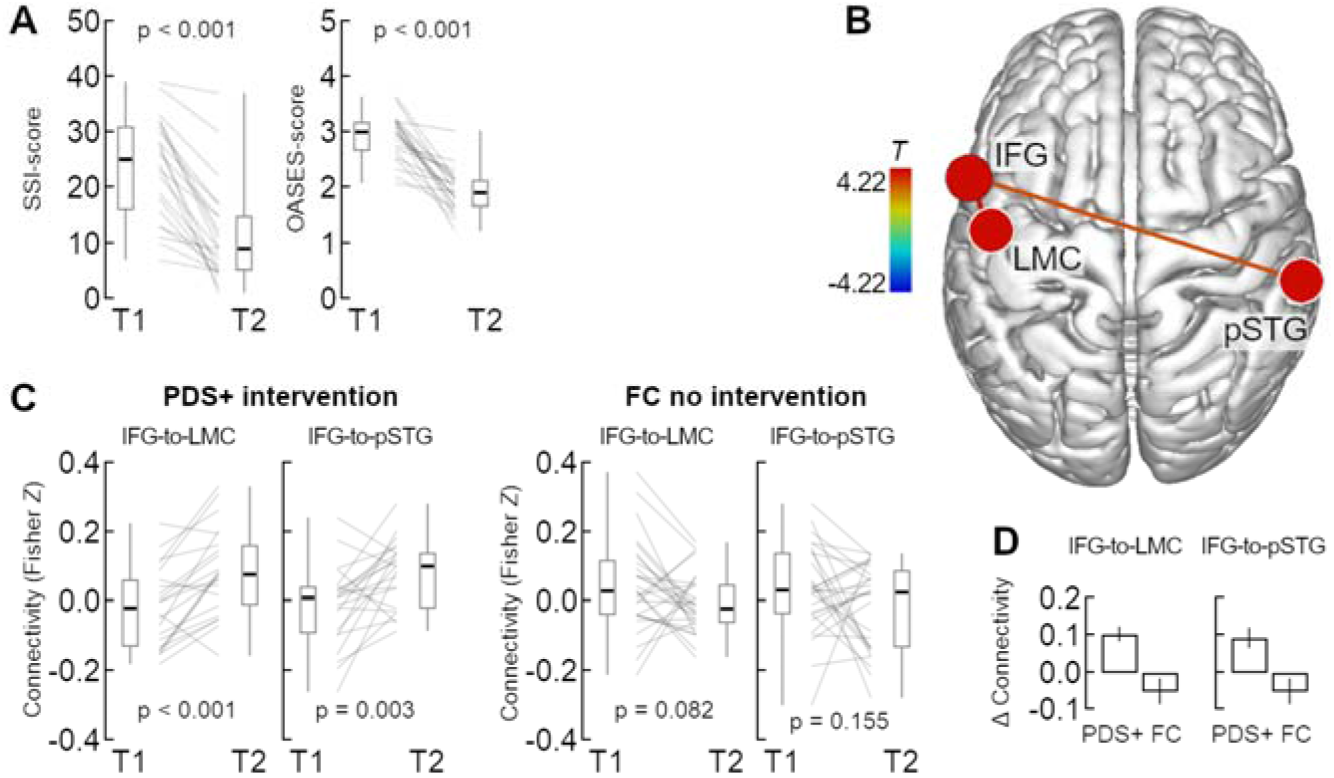
Intervention reduced stuttering and strengthened connectivity. (A) Boxplots display PDS+’ stuttering severity scores pre-(T1) and post-intervention (T2). (B) Rendered brain surface with connections of the Group × Time interaction in the sensorimotor network. (C) Boxplots display rs-fMRI connectivity of PDS+ and fluent controls (FC) at T1 and T2. (D) Barplots display grand mean connectivity changes, i.e. the Fisher Z difference between T2 and T1 (±SEM). IFG = inferior frontal gyrus; LMC = laryngeal motor cortex; pSTG = posterior superior temporal gyrus. Boxplots show whiskers from minimum to maximum, interquartile range, and median values.

**Figure 3.**
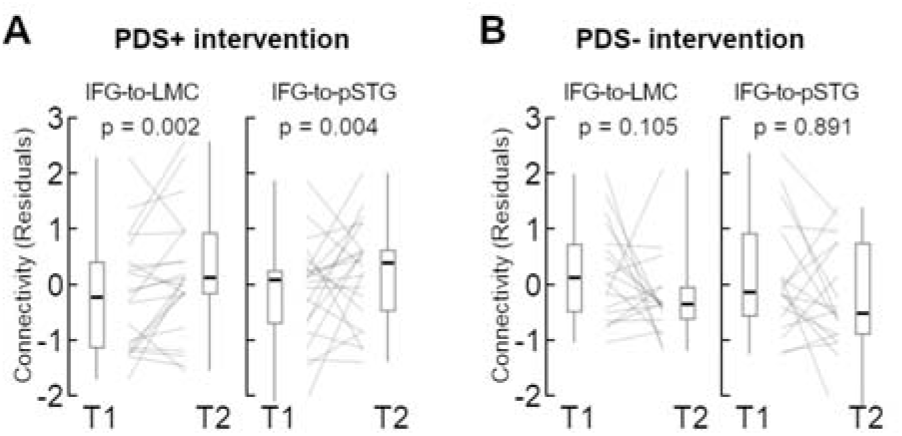
RS-fMRI connectivity corrected for age and stuttering severity (SSI-scores at T1). Boxplots display residuals of (A) PDS+ and (B) PDS− at T1 and T2. Boxplots show whiskers from minimum to maximum, interquartile range, and median values. IFG = inferior frontal gyrus; LMC = laryngeal motor cortex; pSTG = posterior superior temporal gyrus.

### Intervention strengthened sensorimotor network connections

Only one of the four global mixed model ANOVAs revealed significant results. When the sensorimotor integration network seeds were entered in the analysis, there was no effect of Group and no effect of Time, but the left IFG showed a Group × Time × Target interaction with F(10, 39) = 3.46, Intensity = 7.34, *p* = 0.021. Specifically, the interaction was significant for the left IFG-to-left LMC connection with beta = 0.12, T(48) = 4.22, *p* = 0.002, and for the left IFG-to-right pSTG connection, beta = 0.13, T(48) = 3.12, *p* = 0.025 (Fig. 2A). Post-hoc tests showed that connectivity increased in PDS+ but not in FC, and that pre-treatment connectivity was similar between PDS+ and FC, but post-treatment connectivity was greater in PDS+ than FC (Table 3).

### Connectivity changes relate to fluent speaking

To control whether neuroplasticity was related to the intervention and not to stuttering as a condition itself, we tested whether significant changes were only evident in PWS+ or whether a stuttering control group, PDS−, showed similar changes. Because PDS+ and PDS− differed in pre-intervention stuttering severity and age, we fed SSI-4 total scores and age as variables of no interest in the model. The Group × Time interaction was significant for the left IFG-to-LMC connection with beta = 0.17, *T*(34) = 3.42, *p* = 0.002, but not for the left IFG-to-right STG connection with beta = 0.11, T(34) = 1.68, *p* = 0.102. PDS+, but not PDS− showed connectivity increases for both connections (Table 4). Pre- and post-treatment group comparisons were not significant (Table 4).

### No correlation between connectivity changes and fluency enhancement

Neither the change in speech fluency (SSI-4, total score) nor the reappraisal of stuttering (OASES, total score) correlated with the changes in rs-connectivity of PDS+, all p > 0.05.

## Discussion

A one-year, biofeedback-based, speech restructuring training program sustainably facilitated speech fluency of patients with developmental stuttering. Furthermore, neural reorganization included a strengthened synchronization of the IFG pars opercularis with the left LMC and the right pSTG. Thus, resting-state fMRI showed that intensive stuttering intervention remodeled the command-to-execution pathway and the sensory-to-motor pathway within the sensorimotor integration network. Hence, we show here for the first time that a computer-assisted, biofeedback-based, intensive speech training program induced functionally specific, long-term focal changes in task-free brain connectivity.

### Therapy induced a positive shift of brain connectivity

We measured, task-free BOLD fMRI fluctuations with ROI-to-ROI connectivity matrices. Chosen metrics characterize connectivity between pairs of ROIs among predefined sets of regions and yield values varying above and below zero. The metrics called ‘connectivity’ is, at its core, a tanh^−1^-transformed correlation *coefficient*. The naming ‘connectivity’ can be counter-intuitive when a measure with negative values is concerned. We stuck to this nomenclature, following (Biswal et al., 1995; Whitfield-Gabrieli and Nieto-Castanon, 2012). How then can we interpret negative values and changes? The activity waveforms at the two sites of interest underly a number of influences. Some might be direct, between the regions, others indirect. The sum of all influences create the observed correlation, reported as ‘connectivity’. Negative connectivity values signify a negative correlation between the signal waveforms of the two brain regions of interest. This connectivity reflects, in a single number, the net effects of those multiple influences, some driving positive correlations, some driving negative correlations. The overall group median around 0 is consistent with the idea that positively correlating influences and negatively correlating influences are balanced. The fluctuations observed for individual participants, the change of sign from T1 to T2, indicates that the relative magnitude of positive and negatively correlating influences varies. In this framework, the therapy-related shift towards more positive connectivity has to be interpreted as an increase of the influences that drive positive correlations between the two sites. In some participants, this effect occurs relative to a baseline of an overall negative correlation, leading to less negative ‘connectivity’. This shift towards more positively correlating influences on the sensorimotor areas, follows the training and usage of reshaped speech patterns. This is consistent with the hypothesis that the sensorimotor integration necessary for this re(learning) is the cause of this more positively correlating influence.

### Stuttering intervention strengthens connectivity of the left IFG with the left laryngeal motor cortex

The studied intervention encompassed one year of learning and practicing a new speech technique. This speech technique comprises soft voice onsets, consonant lenitions, and controlled sound prolongations (Euler et al., 2009). Thus, voicing and timing are the key features under change throughout the acquisition of the new speech technique. The control of voicing is based on the neural control of the larynx and involves the LMC (Bouchard et al., 2013; Brown et al., 2008; Olthoff et al., 2008; Rödel et al., 2004; Simonyan, 2014; Simonyan et al., 2009), while the control of speech timing involves activity of the posterior part of the inferior frontal gyrus pars opercularis (Clos et al., 2013; Long et al., 2016; Neef et al., 2016). Accordingly, the intensive training incorporated two brain regions, the left LMC and the left IFG, that provide essential neural contributions to fluent speech production.

We found that the intervention strengthened connectivity between the left IFG and the left LMC in the current study. This finding is also consistent with the involvement of the left posterior IFG and left motor cortex in motor learning in general (Papitto et al., 2019) and with the particular involvement of the posterior IFG and the left LMC in speech motor learning (Darainy et al., 2019; Rauschecker et al., 2008). The IFG and the orofacial motor cortex share direct connections (Greenlee et al., 2004) and are commonly co-active under task- and resting-state conditions (Simonyan et al., 2009; Simonyan and Fuertinger, 2015). Furthermore, theories on speech motor control assume the posterior region of the IFG to link the target speech unit to an articulatory code that is subsequently implemented by the motor cortices, which finally orchestrate the articulators, including the larynx (Guenther, 2016). Hence, vital functional connectivity between the left IFG and left LMC is essential for acquiring a new speech technique.

### Stuttering intervention strengthens connectivity of the left IFG with the right posterior superior temporal gyrus

The intervention strengthened co-activity between the left IFG and the right pSTG. The speech motor system has to monitor the auditory feedback signal to correct articulatory errors in natural speech rapidly. Speech-related auditory feedback control involves the right pSTG and task-related co-activations of the left posterior IFG with bilateral pSTGs (Behroozmand et al., 2015; Guenther, 2016; Guenther et al., 2006; Niziolek and Guenther, 2013; Tourville et al., 2008). The DIVA (directions into velocities of articulators) model of speech motor control suggests that the posterior IFG provides feedforward control signals, and the pSTG conveys feedback-based corrective signals (Guenther, 2016). In addition, the right pSTG is associated with spectral auditory feedback control, whereas the left pSTG is more involved during changes in the temporal domain of auditory feedback (Floegel et al., 2020). Learning and practicing a new speech technique addresses neural circuitries of auditory feedback monitoring because patients are constantly required to adjust their speech to fit the new sound pattern of fluency shaping. We suggest that the increased co-activity between left IFG and right pSTG could reflect the frequent recruitment of both brain regions and auditory spectral feedback control mechanisms during learning and practicing the new speech pattern.

Interestingly in our study, PDS+ had no increase of left-hemispheric functional connectivity between the ventral motor cortex vMC and pSTG, contrary to observations in a former study (Kell et al., 2018). This former study used task-related fMRI and showed a reduced left pSTS-to-vMC connectivity before therapy and a strengthened left aSTG-to-vMC connectivity after therapy in PDS+. One possible explanation of why we could not find such an intervention-associated strengthening of left auditory-to-motor coupling could be the seeds’ locations in the current study. While the IFG seeds overlap, MC and STG ROIs are not overlapping. However, another difference between both studies is the time between T1 and T2. The current study investigated the long-term effects. Thus, strengthened connectivity, including the left pSTG could be missing as participants might have shifted auditory feedback control strategies throughout the intervention. One could speculate that temporal features related to the soft production of consonants might be more critical at the early stage of speech motor learning, while spectral features such as vowel length and prosody come later into play.

### Resting-state connectivity is improbable to reflect compensatory brain activity

Here, we measured MRI signal fluctuations in the absence of response demands or external stimulation to describe intervention-induced changes in the speech function-related sensorimotor integration network. It is assumed that spontaneous brain activity at rest relates to the underlying anatomical circuitry (Deco et al., 2013) as supported by diffusion-weighted imaging (Hagmann et al., 2008; Honey et al., 2009). Specifically, it has been suggested that spatially and temporally correlated brain activity at rest arises from neuronal noise between brain areas that share anatomical connections (Deco et al., 2013). In this respect, the current study extends the scope of previous studies where task-related changes in brain activity were observed as a result of the very same intensive stuttering intervention (Kell et al., 2018, 2009; Neumann et al., 2005, 2003). Nevertheless, the current finding of increased sensorimotor learning does not contradict previous conclusions, i.e., normalization of brain activity after intervention (Kell et al., 2018, 2009; Neumann et al., 2005, 2003) as findings are not directly comparable. First, whereas this study investigates long-term effects, the former study tested short-term brain changes directly after the on-site intervention. In addition, the analyses methods of this study highlight learning-related changes and cannot represent neuronal processes related to the occurrence of speech disfluencies. On the other hand, task-related neuroimaging results from learning studies might be confounded by behavioral changes. Using resting-state activity as a neural marker of neuroplasticity rules out that changes in brain activity were induced by changes in task performance (Darainy et al., 2019). In fact, here we provided a purely neurophysiological index of neuroplasticity in the context of an intensive stuttering intervention.

### No correlations between behavioral and connectivity changes

Consistent with our findings, some previous studies reported no correlations between changes in brain activity and changes in speech fluency (Neumann et al., 2003; Toyomura et al., 2015). Others observed correlations between post-treatment speech fluency and task-based brain activity and connectivity (Kell et al., 2018, 2009; Lu et al., 2017) or task-free resting-state connectivity (Lu et al., 2012). These previously reported resting-state functional connectivity changes involved the cerebellum and related the left declive and vermis area to intervention-induced speech fluency (Kell et al., 2009; Lu et al., 2012; Toyomura et al., 2015). In our study, the field of view did not cover the cerebellum, and thus cerebellar ROIs were not included in the analyses. For this reason, it was not possible to test intervention-induced reorganization of the cerebellum.

However, the question still remains of why the significant changes in connectivity reported here do not correlate with speech fluency changes.

### Connectivity changes involve brain hubs of the sensorimotor integration network

We observed no connectivity changes between brain hubs subserving speech planning processes and articulatory convergence. Speech planning processes address working memory resources, and related brain activity is shaped by sequencing demands, syllable complexity, and length (Bohland and Guenther, 2006; Rottschy et al., 2012; Segawa et al., 2015). Articulatory convergence relates to the joint coordination of articulatory movements across the multiple articulatory subsystems (Guenther, 2016). During the intervention, the main focus of practice was controlling the larynx to use soft voice onsets and consonants together with slow speech. This technique requires primarily sensorimotor control and monitoring of the intended auditory target and might unburden or even facilitate speech planning and articulatory convergence. Thus, speech planning processes or the coordination of articulators were not addressed to result in task-free functional connectivity changes. Our observation is in line with a recent neuroimaging study showing that PDS exhibit no deficit in learning to produce novel phoneme sequences (Masapollo et al., 2021); task-related fMRI data showed no difference in brain activity between PDS and controls for the articulation of practiced und novel pseudowords. Thus, sensorimotor learning and feedback processing in the context of voice control may be the main drivers of the neuroplasticity in the current study.

Previous studies also related intervention-associated functional activity and connectivity changes to sensorimotor integration (Kell et al., 2018) and normalized prosody processing (Kell et al., 2018). They suggested that fluency-inducing techniques synchronize a disturbed signal transmission between auditory, speech motor planning, and motor areas (Neumann et al., 2003). Moreover, the main conclusion from these previous task-fMRI studies is that fluency-shaping training normalizes brain activity. This conclusion is based on the observation that pretreatment activity was reduced in the left IFG and increased in the right IFG, but normalized to a level that is comparable to the activity in controls after the intervention (Kell et al., 2018, 2009; Neumann et al., 2003). However, in the present study, pretreatment rs-fMRI connectivity was similar between PDS+ and FC. There was neither altered resting-state connectivity of the left nor of the right IFG. Only post-treatment connectivity was increased between hubs within the set of sensorimotor integration ROIs. Thus, the current findings do not directly support the previous normalization account. Instead, current rs-fMRI data suggest that fluency-shaping training recruits connections that support sensorimotor learning and integration. One could speculate that people who do not stutter would recruit similar connections to learn a new speaking behavior and that intensive therapy addresses a vital brain function. This interpretation is in line with rs-fMRI connectivity changes observed after short-term speech motor adaptation (Darainy 2019).

The increased task-fMRI activity of the right IFG is a signature of persistent developmental stuttering (Belyk et al., 2015; Brown et al., 2005; Budde et al., 2014; Neef et al., 2015). On the one hand, previous studies associate the therapy-associated reduction of this hyperactivity in the right IFG with compensatory mechanisms (Kell et al., 2009; Neumann et al., 2003; Preibisch et al., 2003). On the other hand, previous studies from our group suggest that this hyperactivity might be related to hyperactive action inhibition, which could indicate a pathophysiological mechanism that causes stuttering (Hartwigsen et al., 2019; Neef et al., 2018, 2016). Here we show, that the connectivity between the set of ROIs forming the inhibition network remained unchanged. This observation also supports the view that fluency-shaping training addresses vital sensorimotor learning and integration structures rather than the pathophysiological structures themselves.

### Perspectives concerning other therapeutic approaches to ameliorate stuttering

Common stuttering interventions consist of (1) speech motor interventions partly modifying or entirely reshaping laryngeal, articulatory, or respiratory movements, (2) feedback and technology interventions which use, e.g., delayed auditory feedback to enhance fluency, or visual feedback to support speech motor interventions, (3) behavioral modification interventions, or (4) cognitive interventions improving psychological well-being, self-confidence, and self-conception. The current and previous studies tested neurofunctional correlates of brain reorganization for the first two approaches. However, the neurobiological foundation of an intervention-induced relief from stuttering induced by the other two approaches, such as, for example, the cognitive-behavior intervention (Menzies et al., 2016), would be worth studying.

Of most significant importance are future studies in children with persistent stuttering. Cross-sectional morphological studies with children who stutter imply a primary deficit in left frontal brain hubs of speech motor control (Beal et al., 2013; Chang et al., 2019; Garnett et al., 2018; Koenraads et al., 2020). Moreover, compared to fluent peers, children who stutter exhibited a reduced activation of the left dorsal IFG and the left premotor cortex during overt speech production, as shown by fNIRS (Walsh et al., 2017). However, in particular, rs-fMRI is a feasible approach to extend our knowledge about neuroplasticity related to improved fluency or even recovery in young children because it is not required to engage them in a speech task. A longitudinal rs-fMRI study revealed aberrant network organizations in children who persist in stuttering and in children who recovered from stuttering (Chang et al., 2018). Therefore, a significant future objective is to disentangle intervention-associated neural reorganization and maladaptive changes related to manifestation. A better understanding of conditions that facilitate neurotypical brain functioning in children who stutter could provide the neurobiological foundation of therapeutic strategies that sustainably enhance fluency.

### Limitations

The current study was a non-interventional prospective study. Unlike a clinical trial, data collection and patient participation did not interfere with the choice of treatment, sample collection, procedures, or the treatment itself. Specifically, we collected MRI data without interfering with timing or choice of treatment. It is essential to acknowledge that, like everyone else, persons with stuttering try to get the best out of life, and participating in on-site intensive training for multiple weeks means being away from work, family, and daily obligations. For this reason, the chosen design helped us recruit as many participants as possible. However, this approach resulted in the problem that while PDS+ and FC were comparable for age, PDS+ and PDS− differed for age and stuttering severity. Strikingly, our statistics yielded results that were neither influenced by age nor by stuttering severity at T1. Nevertheless, to enhance comparability between groups of stuttering participants, future studies should match participants according to their stuttering severity.

In addition, the study protocol included a task-related fMRI of covert speaking (not reported here), which was acquired before resting-state. Intervention-related changes in brain activity were evident in the left and right rolandic operculum, the right IFG pars triangularis, the right SMG, the left STG, the left temporal pole, and the left amygdala (Primaßin, 2019). However, none of these regions showed intervention related-changes in task-free resting-state activity. Furthermore, the MRI protocol was kept the same for all participants, and thus, possible carry-over effects would have affected all groups in a similar way. Accordingly, although we cannot entirely exclude carry-over effects, these two aspects make them less likely.

Last, test-retest reliability of metrics of spontaneous BOLD-fMRI fluctuations seems to strongly vary between networks (Noble et al., 2019), regions (Donnelly-Kehoe et al., 2019), and over pairs of regions (Pannunzi et al., 2017). For example, the default mode network and frontal network seem most reliable, and subcortical networks seem least reliable (Noble et al., 2019), which might be related to stronger connectivity estimates in cortical compared to subcortical networks (Noble et al., 2017). Selected sets of seed ROIs were defined based on task-fMRI activity, an accepted approach to evaluate rs-fMRI activity (Hausman et al., 2020; Pando-Naude et al., 2019). This approach was motivated by the presumption that in the absence of a task, brain regions that typically activate together during task performance show strong correlations with one another at rest (Deco et al., 2013, 2011; Vahdat et al., 2011), and the current study was particularly interested in the brain hubs involved in speech motor learning and processing. Because the selected sets of ROIs do not constitute classical resting-state fMRI networks, such as the default network, they seem particularly susceptible to variability and thus to a low test-retest reliability. Reliability across sessions is vital for interpreting longitudinal studies (Birn et al., 2013). The statistical significance of increased connectivity in the face of such variability suggests to us that the effect is of considerable magnitude.

## Conclusion

A one-year practice of fluency-shaping speech techniques boosts the synchrony of spontaneous brain activity in core hubs of speech timing and voice control. Thus, successful speech restructuring shapes sensorimotor integration networks and is reflected in a long-lasting, focal, neurofunctional reorganization.

## Acknowledgments

This work was supported by the DFG (SO 429/4-1 to M.S). We thank Bettina Helten for the co-analysis of the speech samples, Michael Bartl for supporting the organization of the behavioral data, and Britta Perl and Ilona Pfahlert for assistance with the acquisition of the MRI data.

## Author contributions

AK, AP, PD, WP, MS, and NEN conceptualized and designed the study. AP acquired the data. AK and NEN analysed the rs-fMRI data. AK and NEN interpreted the data, drafted, and revised the manuscript for content. All authors reviewed the manuscript.

## Additional Information

The author(s) declare no competing interests.

