## Supplementary Table 1 for "Fluency shaping increases integration of the command-to-execution and the auditory-to-motor pathways in persistent developmental stuttering"

### Supplementary Tables

#### Supplementary Table 1: Demographic information of the individual participants

| ID | Group | Sex | Age, years | Education | Handedness (LQ) | MRI Interval, months | Stuttering onset, years | Years since last treatment |
|---|---|---|---|---|---|---|---|---|
| P_01 | PDS+ | male | 39.58 | 2 | 81.81 | 10 | 6 | 29 |
| P_02 | PDS+ | male | 23.42 | 2 | -100.00 | 10 | 6 | 1 |
| P_03 | PDS+ | male | 29.08 | 3 | 81.81 | 12 | 2 | 19 |
| P_04 | PDS+ | male | 27.83 | 3 | 100.00 | 12 | 4 | 16 |
| P_05 | PDS+ | male | 24.08 | 2 | 100.00 | 12 | 5 | 7 |
| P_06 | PDS+ | male | 30.00 | 2 | 80.00 | 11 | 9 | nan |
| P_07 | PDS+ | male | 15.92 | 2 | -100.00 | 12 | 4 | 1 |
| P_08 | PDS+ | male | 18.50 | 3 | 100.00 | 12 | 5 | nan |
| P_09 | PDS+ | male | 20.75 | 4 | 66.66 | 12 | 3 | 4 |
| P_10 | PDS+ | male | 27.25 | 5 | 100.00 | 11 | 3 | 17 |
| P_11 | PDS+ | female | 17.92 | 3 | 100.00 | 12 | 2 | 2 |
| P_12 | PDS+ | male | 15.42 | 1 | 81.81 | 12 | 3 | 5 |
| P_13 | PDS+ | male | 27.92 | 2 | 100.00 | 12 | 2 | 1 |
| P_14 | PDS+ | female | 15.75 | 1 | 100.00 | 11 | 13 | 2 |
| P_15 | PDS+ | male | 18.58 | 3 | 81.81 | 11 | 4 | 2 |
| P_16 | PDS+ | male | 14.42 | 1 | 100.00 | 12 | 2 | 7 |
| P_17 | PDS+ | male | 15.92 | 1 | 81.81 | 10 | 4 | 8 |
| P_18 | PDS+ | male | 53.67 | 6 | 100.00 | 13 | 12 | 25 |
| P_19 | PDS+ | male | 21.00 | 2 | 100.00 | 11 | 4 | 12 |
| P_20 | PDS+ | male | 31.58 | 6 | 81.81 | 12 | 2 | 3 |
| P_21 | PDS+ | male | 57.00 | 2 | 100.00 | 14 | 6 | 1 |
| P_22 | PDS+ | male | 16.67 | 2 | 66.60 | 10 | 4 | nan |
| P_23 | PDS- | male | 37.50 | 2 | 100.00 | 11 | 2.5 | 29 |
| P_24 | PDS- | male | 28.33 | 6 | 100.00 | 10 | 2 | 2 |

| | | | | | | | | |
|---|---|---|---|---|---|---|---|---|
| P_25 | PDS- | male | 28.17 | 6 | 100.00 | 12 | 6 | 8 |
| P_26 | PDS- | male | 25.58 | 3 | 100.00 | 10 | 2 | 4 |
| P_27 | PDS- | male | 34.17 | 6 | 63.63 | 11 | 2 | 15 |
| P_28 | PDS- | male | 27.92 | 3 | 53.84 | 10 | 6 | 13 |
| P_29 | PDS- | female | 27.50 | 6 | 66.60 | 15 | 3 | 8 |
| P_30 | PDS- | male | 35.58 | 6 | 81.81 | 15 | 3 | 2 |
| P_31 | PDS- | male | 46.75 | 6 | 81.82 | 12 | 2 | 16 |
| P_32 | PDS- | male | 43.33 | 6 | 100.00 | 11 | 3 | 10 |
| P_33 | PDS- | male | 34.25 | 3 | 81.81 | 12 | 2 | nan |
| P_34 | PDS- | male | 30.42 | 6 | 100.00 | 11 | 10 | 11 |
| P_35 | PDS- | male | 46.33 | 7 | 100.00 | 12 | 14 | 14 |
| P_36 | PDS- | male | 43.50 | 7 | 100.00 | 11 | 2 | 15 |
| P_37 | PDS- | male | 34.58 | 2 | -20.00 | 12 | 5 | nan |
| P_38 | PDS- | male | 33.33 | 6 | 66.60 | 11 | 7 | nan |
| P_39 | PDS- | female | 27.17 | 3 | 100.00 | 11 | 10 | 3 |
| P_40 | PDS- | male | 42.50 | 5 | 66.60 | 11 | 8 | 31 |
| P_41 | FC | male | 31.67 | 6 | 81.81 | 11 | | |
| P_42 | FC | male | 34.75 | 2 | 81.81 | 12 | | |
| P_43 | FC | male | 23.50 | 5 | -66.60 | 11 | | |
| P_44 | FC | female | 19.92 | 3 | 100.00 | 11 | | |
| P_45 | FC | male | 27.75 | 3 | 100.00 | 14 | | |
| P_46 | FC | male | 20.25 | 3 | 100.00 | 13 | | |
| P_47 | FC | male | 28.33 | 6 | 100.00 | 12 | | |
| P_48 | FC | male | 27.08 | 3 | 60.00 | 12 | | |
| P_49 | FC | male | 24.08 | 5 | 66.60 | 12 | | |
| P_50 | FC | male | 30.08 | 6 | 66.60 | 12 | | |
| P_51 | FC | male | 20.67 | 3 | 81.81 | 11 | | |
| P_52 | FC | male | 22.42 | 5 | 66.66 | 11 | | |
| P_53 | FC | female | 17.00 | 2 | 100.00 | 12 | | |
| P_54 | FC | male | 30.58 | 6 | 100.00 | 11 | | |
| P_55 | FC | female | 16.83 | 1 | -66.60 | 11 | | |
| P_56 | FC | female | 23.42 | 5 | -54.00 | 11 | | |
| P_57 | FC | male | 27.00 | 4 | 42.80 | 11 | | |
| P_58 | FC | male | 52.42 | 3 | 100.00 | 10 | | |
| P_59 | FC | male | 27.00 | 5 | 100.00 | 11 | | |
| P_60 | FC | male | 14.17 | 1 | 100.00 | 11 | | |
| P_61 | FC | male | 18.67 | 2 | 100.00 | 12 | | |
| P_62 | FC | male | 17.42 | 2 | 81.81 | 11 | | |
| P_63 | FC | male | 17.67 | 2 | 100.00 | 11 | | |

| | | | | | | |
|---|---|---|---|---|---|---|
| P_64 | FC | male | 27.92 | 3 | 100.00 | 11 |
| P_65 | FC | male | 26.50 | 3 | 81.81 | 11 |
| P_66 | FC | male | 25.25 | 5 | 100.00 | 11 |
| P_67 | FC | male | 26.75 | 4 | 100.00 | 11 |
| P_68 | FC | male | 23.08 | 3 | 100.00 | 11 |

*Abbreviations: PDS+: stuttering patients with stuttering intervention; PDS-: stuttering patients without stuttering intervention; FC: fluent controls; achieved education levels were 1 = still attending school, 2 = school, 3 = high school, 4 = <2 years college, 5 = 2 years college, 6 = 4 years college, 7 = postgraduate*

**Supplementary Table 2: Total scores of SSI and OASES at T1 and T2 per participant**

| ID | Group | T1 SSI (totals score) | T2 SSI (total score) | T1 OASES (total score) | T2 OASES (total score) |
|---|---|---|---|---|---|
| P_01 | PDS+ | 28 | 17 | 3.17 | 1.79 |
| P_02 | PDS+ | 16 | 4 | 3.18 | 1.53 |
| P_03 | PDS+ | 23 | 15 | 2.07 | 1.91 |
| P_04 | PDS+ | 27 | 5 | 3.11 | 1.66 |
| P_05 | PDS+ | 38 | 30 | 3.23 | 2.13 |
| P_06 | PDS+ | 13 | 6 | 2.52 | 2.05 |
| P_07 | PDS+ | 22 | 9 | *2.65 | *1.51 |
| P_08 | PDS+ | 13 | 8 | 2.96 | 2.13 |
| P_09 | PDS+ | 19 | 7 | 3.00 | 1.83 |
| P_10 | PDS+ | 10 | 5 | 2.18 | 1.63 |
| P_11 | PDS+ | 18 | 5 | 2.33 | 2.06 |
| P_12 | PDS+ | 39 | 37 | *2.69 | *2.33 |
| P_13 | PDS+ | 30 | 17 | 2.74 | 2.48 |
| P_14 | PDS+ | 7 | 5 | *2.51 | *1.41 |
| P_15 | PDS+ | 24 | 2 | 3.18 | 2.08 |
| P_16 | PDS+ | 31 | 11 | *2.87 | *2.24 |
| P_17 | PDS+ | 26 | 15 | *3.21 | *1.65 |
| P_18 | PDS+ | 32 | 12 | 2.99 | 1.81 |
| P_19 | PDS+ | 28 | 1 | 3.10 | 1.22 |
| P_20 | PDS+ | 32 | 9 | 3.19 | 3.02 |
| P_21 | PDS+ | 33 | 22 | 3.64 | 2.15 |
| P_22 | PDS+ | 11 | 9 | *3.59 | *1.89 |

| | | | | | |
|---|---|---|---|---|---|
| P_23 | PDS- | 28 | 26 | 2.99 | 2.72 |
| P_24 | PDS- | 7 | 8 | 1.38 | 1.27 |
| P_25 | PDS- | 5 | 8 | 1.58 | 1.67 |
| P_26 | PDS- | 15 | 13 | 1.30 | 1.27 |
| P_27 | PDS- | 4 | 4 | 1.89 | 1.87 |
| P_28 | PDS- | 22 | 15 | 2.04 | 1.80 |
| P_29 | PDS- | 14 | 12 | 2.21 | 2.25 |
| P_30 | PDS- | 16 | 18 | 2.13 | 2.45 |
| P_31 | PDS- | 42 | 44 | 2.25 | 2.17 |
| P_32 | PDS- | 7 | 10 | 2.08 | 1.91 |
| P_33 | PDS- | 20 | 22 | 1.73 | 1.71 |
| P_34 | PDS- | 10 | 4 | 2.16 | 2.20 |
| P_35 | PDS- | 11 | 11 | 1.86 | 1.92 |
| P_36 | PDS- | 32 | 35 | 2.46 | 2.28 |
| P_37 | PDS- | 5 | 3 | 1.83 | 1.69 |
| P_38 | PDS- | 14 | 16 | 2.44 | 2.13 |
| P_39 | PDS- | 15 | 16 | 2.10 | 2.12 |
| P_40 | PDS- | 11 | 10 | 1.80 | 1.98 |

*Abbreviations: PDS+: stuttering patients with stuttering intervention; PDS-: stuttering patients without stuttering intervention; SSI: stuttering severity index; OASES: Assessment of the Speaker's Experience of Stuttering; T1: first measurement - prior to intervention; T2: second measurement - after intervention; * OASES – T: Assessment of the Speaker's Experience of Stuttering for Teenagers (Yaruss and Quesal, 2006)*

**Supplementary Table 3: Brain hubs of speech (motor) planning**

| Brain hub – anatomical label | ROI Label | X | Y | Z |
|---|---|---|---|---|
| Inferior frontal gyrus, pars opercularis | L IFG | -56 | 8 | 21 |
| Parietal operculum | L OP | -54 | -22 | 22 |
| Inferior parietal lobe | L IPL | -54 | -36 | 22 |
| Inferior parietal lobe | R IPL | 48 | -56 | 32 |
| Middle temporal gyrus | L MTG | -58 | 0 | -30 |

*All coordinates refer to MNI-space. L = left, R = right. Coordinates were derived from (Neef et al., 2016) and (Clos et al., 2013)*

**Supplementary Table 4: Brain hubs of articulatory convergence**

| Brain hub – anatomical label | ROI Label | X | Y | Z |
|---|---|---|---|---|
| Supplementary motor area | L SMA | -3 | -1 | 57 |
| Supplementary motor area | R SMA | 4 | -6 | 64 |
| Precentral gyrus, | L PrCG | -50 | -4 | 35 |
| Ventral precentral gyrus | R vPrCG | 53 | -7 | 36 |
| Precentral gyrus | R PrCG | 59 | 3 | 19 |
| Postcentral gyrus | L PoCG | -46 | -15 | 15 |
| Putamen | L Put | -25 | -4 | -3 |
| Putamen | R Put | 24 | -2 | 0 |
| Thalamus | L Th | -12 | -16 | 1 |
| Thalamus | R Th | 15 | -15 | 1 |

*All coordinates refer to MNI-space. L = left, R = right. Coordinates were derived from (Guenther, 2016)*

**Supplementary Table 5: Brain hubs of inhibition**

| Brain hub – anatomical label | ROI Label | X | Y | Z |
|---|---|---|---|---|
| Inferior frontal gyrus, pars opercularis | R IFG | 50 | 18 | 6 |
| Supplementary motor area[a] | R SMA | 6 | 18 | 48 |
| Insula[a] | R In | 36 | 18 | 0 |
| rostral cingulate zone[a] (median cingulate paracingulate) | R rCC | 4 | 26 | 38 |
| Subthalamic nucleus[b] | R STN | 8 | -13 | 7 |

*All coordinates refer to MNI-space. L = left, R = right. [a]Coordinates were derived from (Zhang et al., 2017); [b]Coordinates were derived from STN-Atlas Forstmann (Keuken and Forstmann, 2015)*

**Supplementary Table 4: Summary of changes in behavioral outcome measures**

| | Intervention group (n = 22) | | Stuttering controls (n = 18) | | Fluent controls (n = 28) | |
|---|---|---|---|---|---|---|
| | T1 | T2 | T1 | T2 | T1 | T2 |
| SSI-4 total | 25 (14.3) | 9 (10.0) | 14 (11.3) | 13 (9.0) | 0 (4.0) | 0 (2.3) |
| OASES total | 3 (0.5) | 2 (0.5) | 2 (0.4) | 2 (0.5) | – | – |

*Ordinal-scaled variables are presented as median (interquartile range).*
